# Primate Social Organization Evolved from a Flexible Pair-Living Ancestor

**DOI:** 10.1101/2022.08.29.505776

**Authors:** Charlotte-Anaïs Olivier, Jordan S. Martin, Camille Pilisi, Paul Agnani, Cécile Kauffmann, Loren Hayes, Adrian V. Jaeggi, C. Schradin

**Author notes:** Charlotte-Anaïs Olivier, **Email:**. shared first authorship (Charlotte-Anaïs Olivier, Jordan S. Martin). shared last authorship (Adrian V. Jaeggi, C. Schradin). **Author Contributions:** CAO, JSM, AVJ and CS wrote the original draft; CAO, CP, PA, CK and CS collected the data; JSM, AVJ, CAO, CS and CP analyzed data; CAO, JSM, AVJ, CS, LH, and PA reviewed and edited the draft.

## Abstract

Explaining the evolution of primate social organization has been fundamental to understand human sociality and social evolution more broadly. It has often been suggested that the ancestor of all primates was solitary and that other forms of social organization evolved later. However, previous research included the assumption that many understudied primate species were solitary, then finding transitions to more complex social systems being driven by various life history traits and ecological factors. Here we show that when intra-specific variation is accounted for, the ancestral social organization of primates was variable, with the most common social organization being pair-living but with approximatively 15-20% of social units of the ancestral population deviating from this pattern by being solitary living. We built a detailed database from primary field studies quantifying the number of individuals (social units) expressing different social organizations in each population. We used Bayesian phylogenetic models to infer the probability of each social organization, conditional on several socio-ecological predictors, in ancestral populations. Body size and activity patterns had large effects on transitions between types of social organizations. Our results challenge the assumption that ancestral primates were solitary and that pair-living evolved afterwards. Moreover, our results emphasize the importance of focusing on field data and accounting for intra-specific variation. Pair-living is evolutionary ancient, likely caused by reproductive benefits such as access to partners and reduced intra-sexual competition, with more complex social structure (pair-bonding) and care systems (biparental and allo-parental care) evolving later.

**Significance Statement:** Was the ancestor of all primates a solitary-living species? Did more social forms of primate societies evolve from this basic and simple society? The dogma has been that the answer is yes. We used a modern statistical analysis, including variations within species, to show that the ancestral primate social organization was most likely variable. Most lived in pairs, and only 15-20% of individuals were solitary. Living in pairs was likely ancient and caused by reproductive benefits, like access to partners and reduced competition with the sexes. More complex social elaborations like pair-bonds, and biparental and allo-parental care, probably evolved later.

## Introduction

Understanding primate social evolution is central to understand our own social ancestry. Numerous comparative studies have inferred that the ancestor of all primates was nocturnal, small, arboreal and solitary (1–3). Previous research explained transitions from solitary living to more complex social systems by various ecological factors and life history traits (1–5). The inferred ancestral solitary stage hinged largely on prosimians, which are basal to the primate phylogeny but understudied and previously often assumed to be solitary living (6). However, several field studies over the last decades indicate prosimians to be more social (1, 6) and often pair-living (7).

Social systems are composed of different components including the social organization (composition of social units), social structure (interactions between individuals), care system (who cares for infants), and mating system (who mates with who) (8, 9). It has been argued that these components should be studied independently from each other to understand social evolution, especially in primates (1, 10, 11). For example, pair-living as a form of social organization has often been equated with monogamy, but monogamy refers to a mating system (1, 11–14). Importantly, pair-living species can vary significantly in their mating system, i.e. the degree of extra-pair paternity (15, 16). Similarly, primate social organization varies greatly between (1, 3) but also within species (6, 17). Previous studies were statistically limited by assigning a single type of social organization to each species, such that the analysis could only consider between but not within species variation (1–3, 18).

Here we examined whether taking intra-specific variation in social organization (IVSO) into account and focusing on data from field studies, including many recent studies on nocturnal prosimians, but excluding assumptions about non-studied species, changes our estimate of the ancestral primate social organization. Primate social evolution is assumed to largely depend on ecology and life history (19, 20). For example, small species are generally assumed to be solitary and large species to be group-living, with group composition depending on diet (3, 8, 19). Species living in heterogeneous habitat are predicted to have a more variable social organization (21). Therefore, we tested in how far multiple ecological and life history factors influenced primate social organisation (Table S2).

We assembled a database on the social organization of 499 populations of 216 primate species observed in the field, as published in the primary peer-reviewed literature. Rather than selecting a single social organization per species we treated each study population as the unit of analysis. Furthermore, within each population we counted the number of social units exhibiting different social organizations, allowing us to quantify within population variation (Fig. 1). Therefore, our statistical approach allowed us to consider variation in social organization (i) between species, (ii) between populations of species and, (iii) between social units within populations. We developed a flexible Bayesian phylogenetic GLMM framework to partition this extensive variation in social organization across populations, species, and superfamilies, as well as to infer its phylogenetic and socio-ecological determinants. Using a multinomial likelihood, we modelled the relative frequency of each social organization being observed within each population, adjusting for phylogeny and research effort. We defined the ‘primary social organization’ as the social organization with the greatest probability of being observed within a population. As a second response variable in the same model, we used a binomial likelihood to directly account for the degree of intrapopulation variation in social organization (IVSO) observed in each population, calculated as the proportion of social units deviating from the most frequent social organization (Fig. 1). This allowed us to estimate effects of socio-ecology and phylogeny on IVSO per se, irrespective of the relative probabilities of specific social organizations within a population.

**Fig 1.**
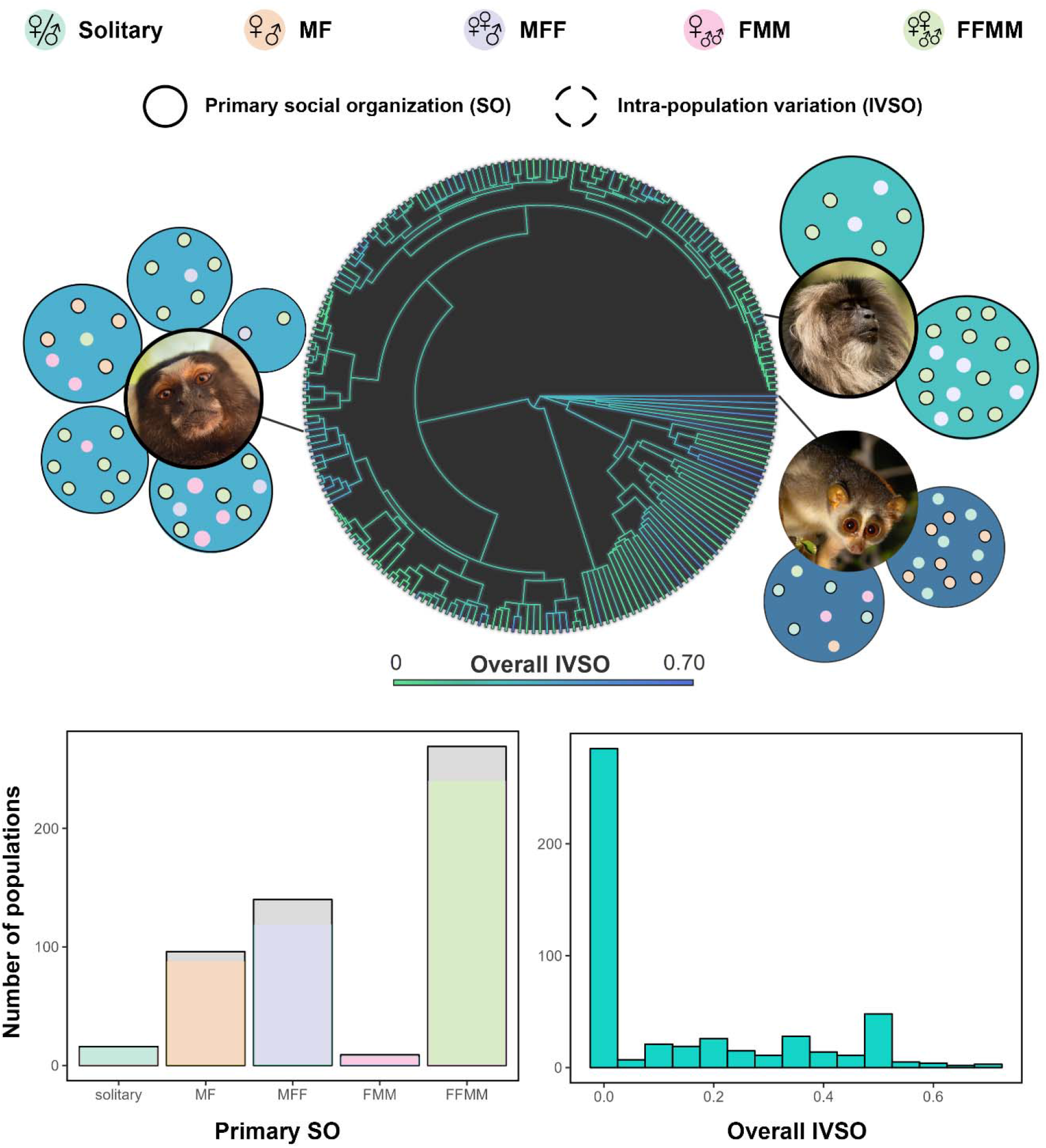
The distribution of social organizations across extant primate populations. Three examples taken from field research on the slender loris (*Loris lydekkerianus*), common marmoset (*Callithrix jacchus*), and lion-tailed macaque (*Macaca silenus*) are shown. The top panel demonstrates how we coded social organization per population as solitary, male-female (MF) or pair living, single male multi-female (MFF), single female multi-male (FMM), or multi-male and multi-female (MMFF). Large circles (middle) around pictures represent different populations of the species. Smaller circles within each large circle represents a single social unit within a population, with color correpsonding to the social organization observed. The phylogeny reflects a simple contour mapping of overall IVSO (# units deviating from primary social organization / total # units) across taxa in our database. Note that the branch lengths have been arbitrarily modified for visual clarity and should not be directly interpreted. The low panel shows the total number of populations in our dataset exhibiting each form of primary social organization, as well as overall IVSO (binwidth = 0.05). Gray bars represent uncertainty in the primary social organization for populations exhibiting two social organization with equally high frequency.

## Results

### Distribution of social organization in extant primates

We observed relatively high rates of pair-living (primary social organization in 26% of populations; MF) and low rates of solitary organization (primary social organization in 3% of populations) in our database (Fig. 1). Previous studies (1, 3, 5) estimated many more species to be solitary living, as they were classified most solitary foragers as also having a solitary social organization. Our database challenges this assumption, showing that among 20 solitary foraging populations (from 15 species) only 8 had solitary living as primary form of social organization, while in the remaining 12 populations pairs or groups shared one home range. The most common form of social organization in extant primates are multi-male multi female groups, followed by one male multiple female groups and pairs, while both one female multiple-male groups and solitary populations were rare (Fig. 1). Many species (47%) and populations (43%) exhibited more than one social organization, demonstrating that primate populations show substantial levels of variation in social organization (Fig. 1).

### Variance due to phylogeny, ecology and life history

We first wanted to know how much variation in primate social organization and IVSO is explained by phylogenetic history, current ecological and life history conditions, or unmeasured effects at the levels of populations, species, and superfamilies. Ecological and life history conditions included habitat heterogeneity, open vs closed habitats, foraging strategy, locomotion, activity pattern, body size, and dietary reliance on fruit, foliage, seeds, or animal protein. Using multiple imputation to leverage all predictors despite missing data (see supplementary materials and supplementary data), we found that ecological and life history variables collectively explained only a small-to-moderate proportion of variation in social organization and IVSO (median R2 range: 0.04 – 0.30; Fig. 2A; see supplementary materials for details on the direct and total effects of each predictor with and without imputation). Phylogeny explained a moderate to large proportion of variation in social organization (median λ range: 0.26 – 0.69), although single-female multi-male social organization had much lower phylogenetic signal (median λ = 0.06). Species- or superfamily-level effects, independent of ecology and phylogeny, were weak (median R2 range: 0.01 – 0.13), suggesting against grade shifts. However, population-level heterogeneity was consistently larger (median R2 range: 0.13 – 0.63), indicating a sizable portion of unexplained variation among populations within the same species. These results suggest that while some forms of social organization are conserved within primate lineages, social organization often shows substantial variation among populations that remains unexplained by phylogeny, ecology or life history.

**Fig. 2.**
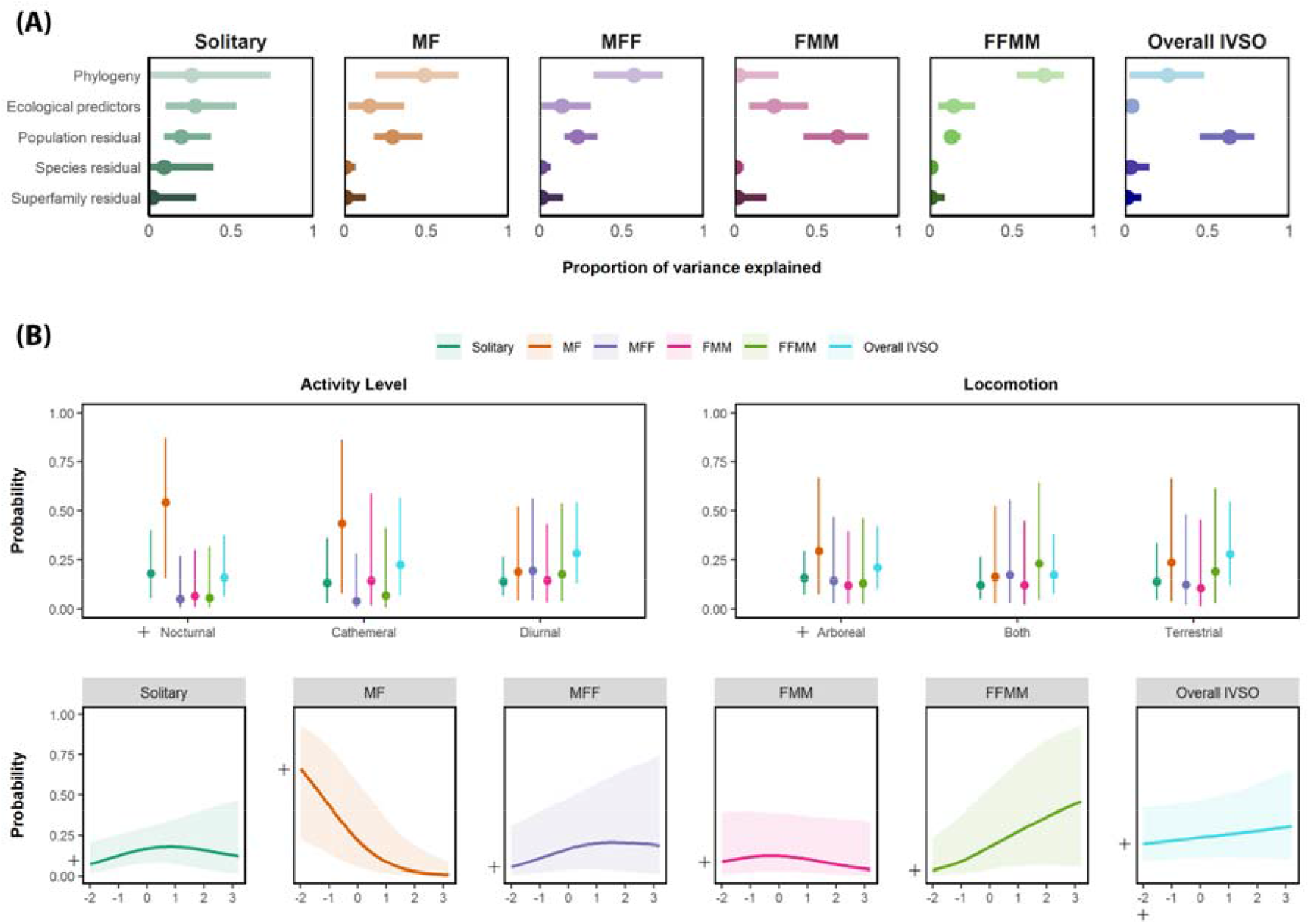
Variation in social organization among extant primates. (**A**) The proportion of variation in each social organization and overall IVSO accounted for by phylogenetic history, ecological and life-history factors (“ecological predictors”: habitat type and heterogeneity, diet, foraging style, locomotion, activity pattern, and body size), research effort (number of published studies on a population, centered within superfamilies), as well as remaining residual (unexplained) variation among populations, species, and superfamilies. Dots indicate posterior medians and lines indicate 90% Bayesian credible intervals (CIs). (**B**) Total effects of the ecological and life history factors (activity level, locomotion, and body size) used to predict ancestral social organization and IVSO. Cross lines (**+**) indicate assumed states used for ancestral state prediction (i.e. nocturnal activity, arboreal locomotion, and low [−2 SD] body size. Thick lines indicate posterior medians and ribbons indicate 90% Bayesian CIs.

### Ecological predictors and reconstruction of the ancestral state

Next, we estimated the probability of each social organization and degree of IVSO for the last common ancestor of all primates, contingent on predictors. Specifically, current evidence strongly suggest that ancestral primates were small-bodied, arboreal, and nocturnal (22, 23), hence body size, locomotion and activity pattern can be used to make more informed inferences about ancestral states. Conversely, since the literature does not offer strong a priori expectations about ancestral habitat types, foraging strategies, or specific dietary patterns we excluded these variables from this model, which had the added benefit of not requiring imputation.

The total effects of body size, locomotion and activity pattern on social organization are shown in Fig. 2B. Overall, pair-living was more common with nocturnal than diurnal activity (median Δ probability = 0.30, 90% CI [0.04, 0.58]). Pair-living was also more likely among smaller bodied species (−1SD) compared to average-sized species (median Δ probability = 0.19, 90% CI [0.05, 0.35]) or larger species (+1 SD; median Δ probability = 0.32, 90% CI [0.09, 0.59]). Multi-male, multi-female groups were in turn more likely among larger-bodied species compared to average (median Δ probability = 0.08, 90% CI [−0.01, 0.26]) or small species (median Δ probability = 0.16, 90% CI [0.00, 0.47]). No clear effects were observed for differences in locomotion on the probability of social organization, and none of these ecological predictors consistently explained variation in the overall proportion of IVSO across populations.

The oldest known primates were very small (22, 24), and we therefore assumed an ancestral body size of 30g (−2 SD relative to extant species; Fig 2B), which is an upper limit based on current fossil evidence (*19*). We also assumed nocturnal activity and arboreal locomotion. Under these assumptions, pair-living units are the most likely ancestral social organization (median probability = 0.80, 90% CI [0.33, 0.98]), compared to solitary (median Δ probability = 0.72, 90% CI = 0.16 – 0.95) and all forms of group living (all median Δ probabilities ≥ 0.75 and 90% CIs exclude zero; Fig. 3). In addition, there is support for a small proportion of solitary social units occurring in ancestral populations (median probability = 0.07, 90% CI [0.01, 0.26]), while little to no support is provided for the presence of group-living units (lower 90% CIs < 0.01). Put differently, if we could sample 10 social units in an ancestral population, we would expect ~80-90% of those units to be pair-living, but also ~10-20% to be solitary. This pattern is supported by the overall IVSO model, which estimates that approximately 10-20% of social units (median probability = 0.15, 90% CI [0.06, 0.35]) should deviate from the primary social organization (pair-living). Thus, the last common ancestor of all primates most likely had a variable social organization, with most individuals living in pairs but some being solitary. Predicted ancestral IVSOs are also relatively constant across each of the six major primate superfamilies (Fig. 4), suggesting that specific clades do not differ in their propensity for intrapopulation variation. Despite pair living likely being ancestral in early primates, support for pair living is only found for ancestral strepsirrhines (lemurs and lorises) and tarsiers (‘prosimians’). Ancestral cercopithecoids (‘Old World’ monkeys) and ceboids (‘New World’ monkeys) were instead more likely to be group living, suggesting that pair-living is derived in Simiiformes and has evolved secondarily within these two superfamilies.

**Fig. 3.**
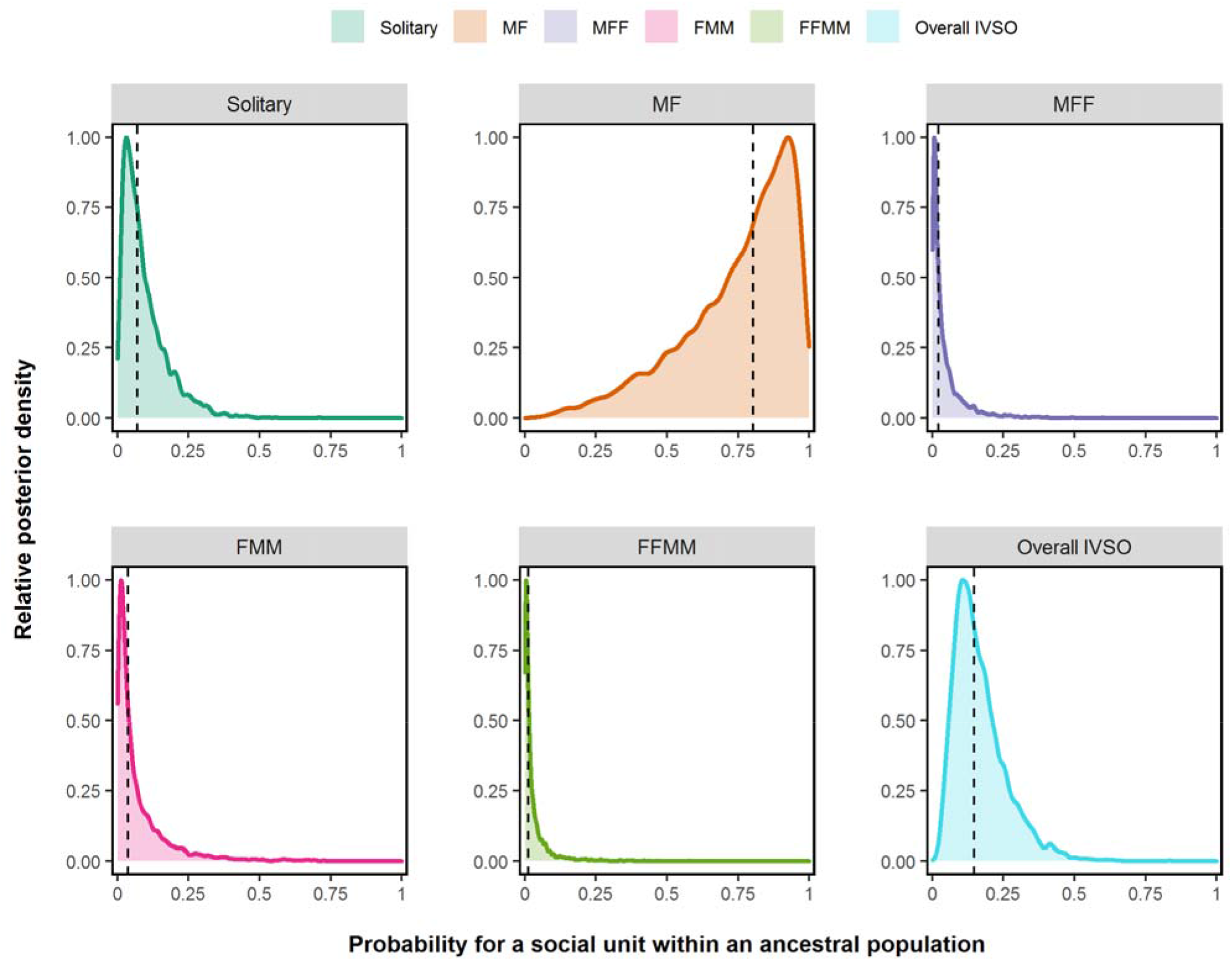
Ecologically informed predictions of social organization in ancestral primate populations. Predicted probabilities of a social unit exhibiting each social organization and some form of IVSO within ancestral populations, assuming the ecological conditions marked in Fig. 2B (nocturnal, arboreal and small), as well as average within-superfamily sampling effort. Scaled posterior densities are shown, with posterior medians indicated by the dotted line.

**Fig. 4.**
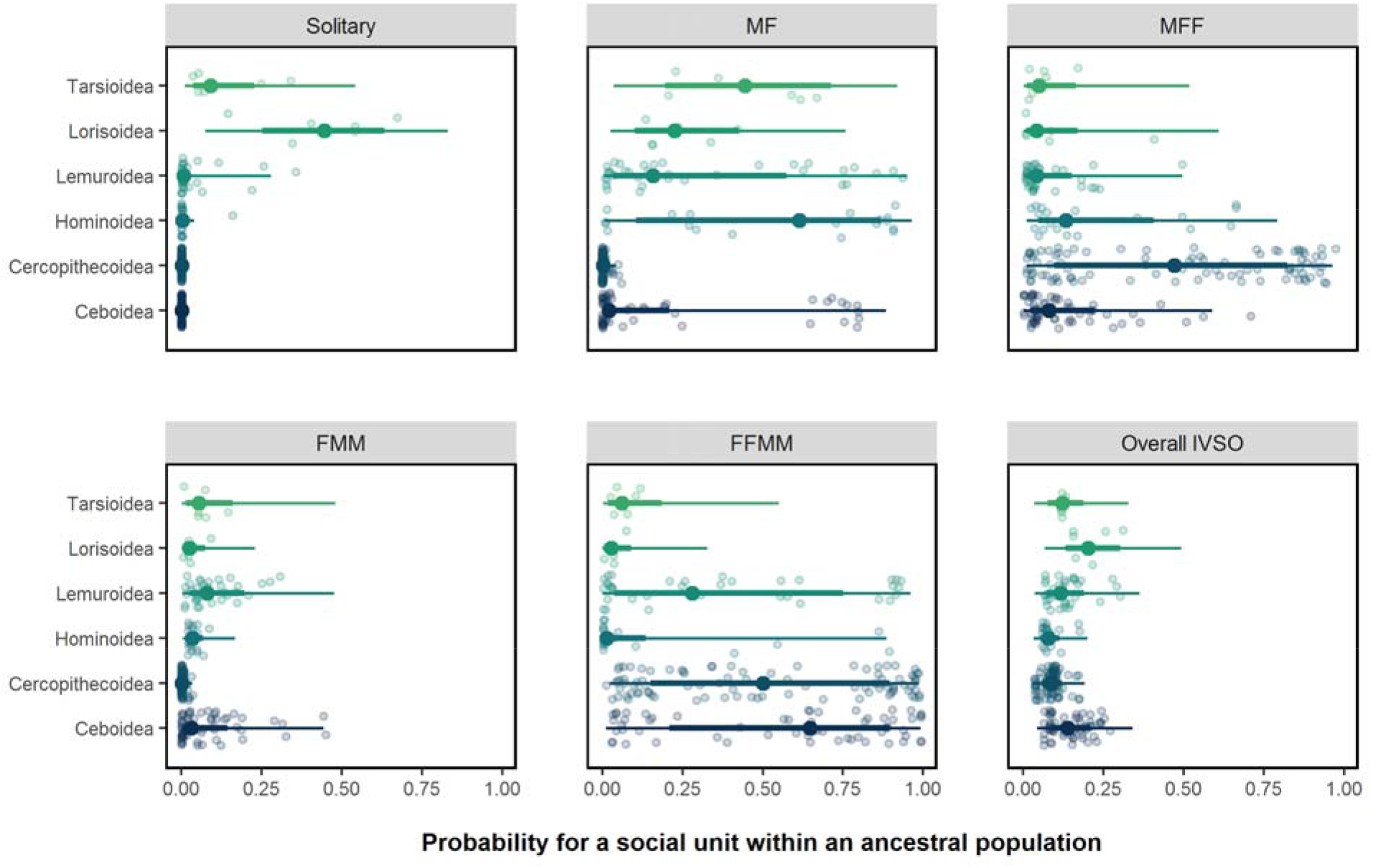
Predictions of ancestral social organization for the six primate superfamilies. The dark dots and lines indicate median probabilities +/- 50% (thick line) and 90% CIs (thin line) for the expected probability of each SO and IVSO in an average population of each primate superfamily. Light circles in turn indicate median predictions for specific species within each superfamily. The root/basal predictions for each clade are the average of the posterior phylogenetically adjusted species predictions, assuming average within-superfamily sampling effort.

## Discussion

Like previous analyses (1, 3, 5), our results suggest that group-living evolved late in primates’ evolutionary history. However, our finding that pair-living was the ancestral primate social organization contrasts with previous studies which found solitary living to be ancestral (1, 3, 5). This difference is not likely a consequence of us having underestimated the occurrence of solitary living in extant primates. Instead, it is likely to be a consequence of our focus on well-studied species and the exclusion of non-studied nocturnal, cryptic species that have often been assumed to be solitary. Multiple field studies revealed that few nocturnal prosimians exhibit solitary living, with instead pair-living being common in these species (7, 25). Accordingly, we found pair-living to be more common in nocturnal prosimians than in diurnal simians, which contrasts previous believes. Further, we focused on the composition of social units (social organization), while most previous studies aimed at explaining “social monogamy”, a concept combining the social organization of pair-living with the social structure of pair-bonding, biparental care and a primarily monogamous mating system (3, 5). This demonstrates how differences in classifying social systems can influence the interpretation of social evolution.

Here, we showed for the first time that the ancestral social organization of primates was variable, with approximately 15% of the individuals in the population deviating from pair-living. Our analysis differs from previous studies by taking IVSO into account. Both Shultz et al 2011 and Lukas & Clutton-Brock 2013 categorised each species as having one social system. Kappeler & Pozzi 2019 were the first to consider IVSO descriptively, but their statistical analysis relied on categorising species into a single form of social organization. Variation is needed for evolutionary change, and it is therefore important to develop statistical tools that take this variation into account. Considering IVSO on the population level allowed us to come to a more realistic estimate of the ancestral social organization of primates.

Social monogamy has often been regarded as a derived form of social system needing specific explanation (1, 3, 5). Thus, it might seem surprising that we found pair-living to be the most likely ancestral social organization. However, pair-living has also been suggested to be the ancestral form in other mammalian orders when considering IVSO, including Artiodactyla (26), Eulipotyphla (27), and in Macroscelidea (28). Moreover, Kappeler & Pozzi 2019 suggested that pair-living is not a derived complex social organization in primates, but an ancestral form before the evolution of more complex social groups.

Our results indicate that pair-living is ancestral in primates, despite other components of “social monogamy” being derived traits present in only a few lineages, including a pair bonding social structure and biparental care system (Fig. 5). Pair-living without pair-bonding as observed in some extant prosimians represents the ancestral primate state. However, our results also show that the direct ancestors of New World and Old World monkeys were most likely not pair-living (Fig. 4). In these taxa, pair-living might have evolved secondarily, and afterwards other components of the social system can have co-evolved. First pair-bonding, as observed in three of the four forms of pair-living social systems, while paternal care only evolved in some New World monkeys and siamangs (Fig. 5). Cooperative breeding with non-breeding helpers only evolved in the callitrichids. Thus, pair-living as in extant primate species can be part of four different social systems (Fig. 5), highlighting the importance to differentiate between different components of social systems (9–11).

**Fig. 5.**
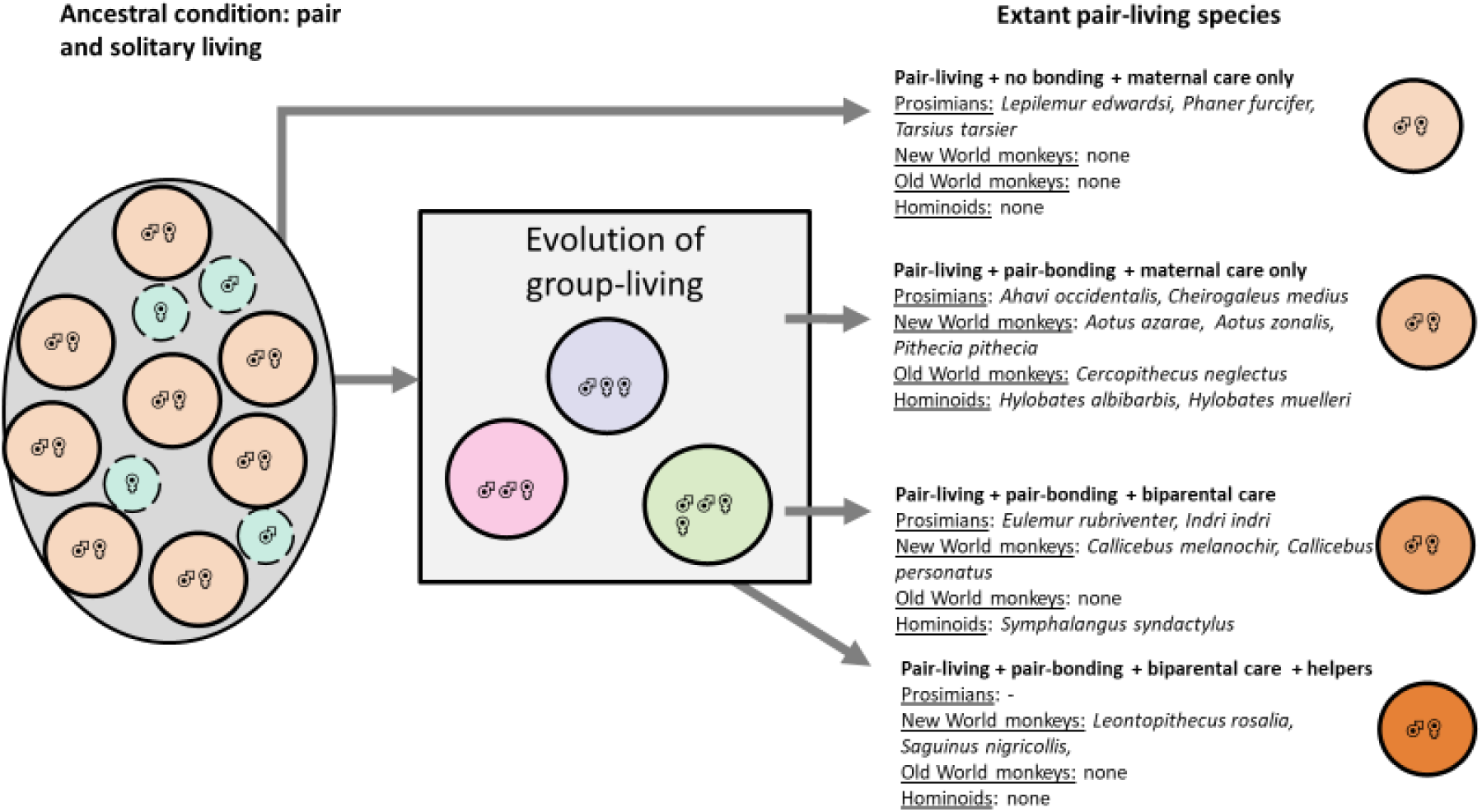
Pair-living in extant primate species and other components of social monogamy. While pair-living is an ancestral state in primates, pair-bonding and paternal care are considered to be independent traits that evolved later and not in all pair-living lineages. Ancestral pair-living seems to be maintained in prosimians (Tarsioidea, Lorisoidea and Lemuroidea), but probably evolved secondarily in the other primate lineages. The large circle represents the ancestral population. Smaller circles represent social units with its outline corresponding to whether this social organization is the most frequently observed or is a form of intra-population variation in social organization (IVSO, dotted line). Colors and symbols represent the different types of social organization as in Fig. 1. Arrows indicate different possible evolutionary pathways. Far right: Pair-living (social organization) occurs in different combinations with the other factors of social systems (right: social structure and care-system), indicating that pair-living can be one component in four different types of social systems (different grades of orange). Examples of pair-living species in the different taxa are shown on the right.

## Materials and Methods

### Materials

#### Definition of social organization

Animal social systems can be characterized by variation in four inter-related components (9): social organization (composition and size of groups), social structure (social interactions), mating system (who mates and who reproduces with whom), and care system (who takes care of the offspring). Previous research has often used heterogenous terminology to describe social systems across taxa, resulting in ambiguous definitions and confounding of distinct selection pressures (29). Here we strictly focused on the composition of groups, a central aspect of social organization. Data for group composition were taken from the methods and results section of peer-reviewed primary literature, avoiding any interpretations from the authors in the discussion. We defined social organization as solitary, pairs (MF), single male + multi-female (MFF), multi-male + single female (FMM), multi-male + multi-female (FFMM), and sex specific, where groups consisted of one sex only.

#### Database on social organization

We identified 450 species of primates using the IUCN (International Union for Conservation of Nature) database (2019). We then conducted literature searches on social organization in Web of Science (Thomson Reuters) and in Google Scholar between January 2016 and September 2019. For each species, we initially search the Latin name of the species and the term “social” (e.g. “*Alouatta caraya* AND social”; “*Gorilla gorilla* AND social”). If no literature on social organization was found, only the Latin name was searched. For several species the Latin name changed over the years, in which case we repeated the search with the previous versions of their names.

Searches on Web of Science were restricted to articles within one of the following three categories: “behavioral science”, “zoology” and “environmental science/ecology”. Only peer-reviewed literature from field studies was considered while reviews, laboratory-based studies and captive studies were ignored, to ensure that the social organization observed by a given species also occurred in their natural habitat (e.g. many species can be kept in captivity in pairs, but this does not mean that pairs occur in nature). For each study, we read the abstract, examined all figures and tables, and searched the following keywords throughout the papers: “social”, “solitary”, “group”, and “pair”. As such, our literature search focused on the data reported in methods and results sections of the peer-reviewed studies, not on the interpretation of the authors regarding the social organization in the introduction or discussion. This search yielded more than 2000 articles that were scanned for information on social organization. Of these, a total of 946 papers contained useable data (83 for Strepsirrhines, 247 for New World monkeys, and 636 for Old World monkeys). Overall, data on social organization were found for 499 populations from 223 species. To determine the forms of social organization present in each population, we recorded the adult sex composition of all social units in a population using the classical definitions from Kappeler and van Schaik (8) and adding “sex specific group” as an additional category (Table S1). Studies that did not report the sex of individuals were not taken into account, following (30).

From each paper, we recorded how often each category of social units was observed e.g. how many solitary individuals, pairs or different groups were recorded. We only recorded solitary living as a form of social organization when both sexes had been observed to be solitary, as single individuals of one sex may often represent dispersers. Whenever individuals were explicitly reported to be dispersers, they were not considered in the recording of social organization. Therefore, to record one social unit of solitary living, at least one solitary male and one solitary female were needed, to make the classification comparable to the criterion used for pair-living (one pair also consists of one male and one female). For example, when 5 solitary males and 4 solitary females were reported, we recorded 4 solitary social units. The same procedure was done with sex-specific groups. We only recorded sex specific groups as a form of social organization when both sexes had been observed to live in unisex groups. To record one social unit of sex-specific groups, at least one group of males and one group of females were needed. For example, when 10 groups of males and 4 groups of females were reported, we recorded 4 sex-specific groups social units. Overall, we only observed sex-specific groups in nine species, indicating that intersex units are the dominant form of social organization in group-living primates. This low count prevented us from drawing meaningful inferences about phylogenetic and ecological effects on the probability of sex-specific units occurring, and we therefore excluded these units from our analyses.

Data were collected at the population level by recording the total number of papers reporting a given social organization in a population. When the same observed individuals and their social units were included in more than one published paper, we considered only the most precise paper, e.g. papers where the precise number of social organizations and/or the sex of individuals was described, to avoid considering the social units of a social organization several times. The total number of studies reporting social organization per population was then recorded in the database to account for any effects of research effort. For example, populations with multiple studies over decades might be more likely to show variation in social organization than populations with only one single study. Similarly, taxa exhibiting greater (or lesser) variation in social organization may be more or less likely to be investigated by researchers.

The database records for each population whether multi-level societies or fission-fusion societies occurred. When multilevel societies (31) occurred within a population, indicating hierarchically structured social organization, we only recorded the composition of the core group defined in the study. We did this because the different core groups within primates’ multilevel societies tend to maintain their social organization across interactions, such as the maintenance of the one male multiple female groups composing large multilevel hamadryas baboon (*Papio hamadryas*) societies (32). In contrast, when fission-fusion societies (33) occurred in the population, suggesting a more fluid social organization, we recorded all forms of group composition observed, as this indicated that individuals of this population could exhibit multiple forms of social organization within their society over time.

#### Intrapopulation variation in social organization (IVSO)

IVSO was identified when different forms of social organization occurred within a population, indicating some degree of behavioral plasticity in a species’ social organization. However, the following cases were not regarded as IVSO: when only one sex had two or more forms of social organization or cases of dispersing individuals (solitary individuals of one sex only) or of alternative reproductive tactics (for example male followers during the breeding season). Environmental disruption such as the death of a dominant breeder or predation of group-members can also change the social organization of a unit (Schradin 2013), but these changes do not reflect the evolved behavioral plasticity we want to explain. Thus, such environmental disruption events were not considered in our database but were recorded separately.

Previous studies have treated IVSO as a distinct category of social organization (26, 28). However, given the wealth of data available for primate social organization, we were able to continuously measure IVSO at the population level as the proportion of social units deviating from the most frequently observed (‘primary’) social organization within a population. In the present study, we therefore conceptualized and measured IVSO as a distinct trait capturing the overall degree of variation in social organization, which may coevolve with the composition and frequency of specific social organizations within a population. This avoided the use of arbitrary thresholds for categorizing the presence or absence of IVSO, retaining the continuous information provided by previous literature, and allowed us to consider how the evolution of specific forms of social organization and the overall degree of IVSO are related across species and populations.

#### Predictors of social organization

We included the following predictors in our Bayesian model to account for potential social, ecological, life-history and methodological causes of variation in social organization: habitat heterogeneity, habitat type, body mass, diet, activity pattern, locomotion, number of studies per population, and foraging strategy (Table S2). Habitat type was recorded from the primary literature and categorized on IUCN classification and used to calculate habitat heterogeneity (total number of habitats per population). Further, we classified the different types of habitat as open, closed or open and closed. Populations were also categorized as having group- or solitary foraging individual or both, depending on information in the primary literature use to categories social organization.

We used two published databases (34, 35) for the predictors body mass, diet, activity pattern and locomotion. First, we compared the two databases to see if their information was very similar. This was the case for locomotion (terrestrial, arboreal and both) and activity pattern (diurnal, nocturnal and cathemeral). However, for body mass and diet we found differences between the two databases, which could undesirably influence our statistical inferences. Body mass was recorded for males and females, with some studies only reporting average body mass across sexes. If for one species more than one measurement for body mass was available for either sex, then we calculated the mean value and the standard deviation. For food, information was available at the species level for average diet composition, including percentage of fruits and foliage (addition of mature leaves, undefined leaves and young leaves) as well as the percentage of seed and animal protein consumed. We took multiple steps to ensure that estimated effects for these variables were robust to variation in diet and body size results between the databases. Whenever there was a difference between the two databases, we checked whether their information was based on the same or on different published studies. If both databases reported the same primary study, we checked the publication ourselves and only utilized the data reported directly in the paper. If the two databases were not based on the same study, we then entered the average result across databases to account for potential heterogeneity and/or measurement error within taxa. We also conducted analyses of body size separately within each database to ensure that aggregated estimates were robust across datasets. No meaningful differences were observed between the databases in the main effects of body size on social organization and IVSO (all Δ*β* 90% CIs included zero), so we used the average species body size between databases for all reported analyses. Information on diet was also combined between the databases when possible to increase sample size, due to heterogeneous patterns of missing data across species.

### Methods

#### Statistical analyses

We developed multilevel phylogenetic models to investigate the evolution of social organization and IVSO across primates, and we conducted all analyses within a flexible Bayesian framework to account for non-Gaussian outcome measures, measurement error in social and ecological data within taxa, as well as uncertainty in phylogenetic relationships across taxa (36–38). Our analyses relied on one of the most recent and up-to-date mammalian phylogenies from the VertLife project (39), ensuring that evolutionary relationships were accurately represented between all species in our dataset. Primary social organization was modelled as a multinomial response variable, appropriate for repeatedly measured categorical data (40), while IVSO was treated as a binomial response, representing the number of groups deviating from the primary social organization out of the total number of groups observed (i.e. each social group observed in a population was coded as either being ‘primary’ = 0 or ‘non-primary’ = 1 in organization). Multi-response models were estimated to simultaneously assess phylogenetic and ecological effects on these measures, as well as to conduct robust ancestral state reconstruction of the primary social organization and magnitude of IVSO expected in basal primate populations. We took multiple steps to integrate uncertainty in our phylogeny and empirical measures during these analyses, which were supported by robustness checks to ensure appropriate inferential stability among models. Conservative, weakly regularizing priors were also used to introduce more realistic assumptions into the estimators, as well as to reduce the risks of inferential bias caused by multiple testing and measurement and sampling error (38, 41). Code for all analyses described in the text can be found at https://github.com/Jordan-Scott-Martin/primate-SO-analysis along with the original database at https://github.com/CharlotteAnaisOLIVIER/Social-organization-of-primates.

For all analyses, we estimated two generalized multilevel phylogenetic models, one for describing the probability of each social organization with a multinomial distribution and the other for describing overall IVSO with a binomial distribution. For population *p* for species *s*, the multinomial model predicted the number of units observed in category *i* as a function of the total number of units observed *n_ps_* and a vector of parameters *θ_ps_* for the relative probabilities of each category compared to the base category, which in this case is solitary social organization.

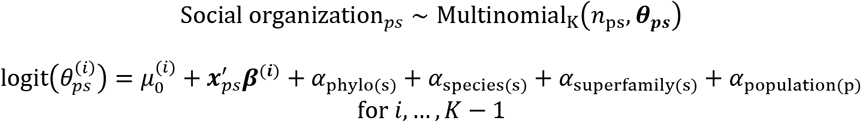

The parameters 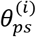 for each category *i* (MF, MFF, FMM, FFMM) as compared to *K* (Solitary) were predicted on the transformed logit scale by a category-specific intercept 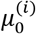, fixed effects ***β^(i)^*** (research effort and ecological predictors), where 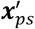 is the transposed vector of population- and/or species-specific predictors, and by the random effects α, which capture Brownian Motion phylogenetic effects *α*_phylo(s)_ as well as any deviations from these phylogenetic predictions at the superfamily, species, and population level. Note that *α*_population(p)_ is as an observation-level random effect capturing overdispersion from the expected variance.

By adjusting for any species-level effects, the *K*-1 intercepts ***θ_ps_*** provide appropriate relative probabilities of non-solitary compared to solitary social organization for an average ancestral population. These values can be transformed to the absolute scale using the logistic function, which facilitates calculating the probability of social units in an average ancestral population showing each of the *K* social organizations. In particular, for solitary and any other social organization *i*

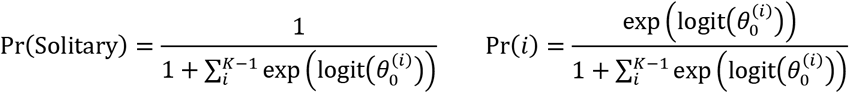

Note that this standard parameterization of the multinomial model can be equivalently specified with *K* intercepts, where 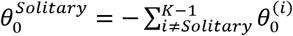. This approach allows for modelling predictors directly on the probability of each category, as shown in Figure 2A, and can be implemented manually in Stan.

The variance explained in social organization by each set of effects can be calculated on the transformed scale using model predictions for the fitted data. Specifically, the total latent variance for social organization *i* is given by

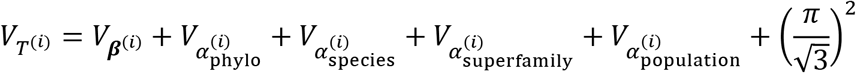

where 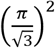 is the theoretical variance of the logit scale. The latent variance explained (*R^2^*), also known as the repeatability or phylogenetic signal λ) can then be estimated by

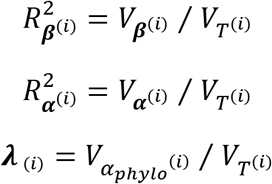

The same approach is taken for predicting the total probability *τ_ps_* of IVSO given the number of social units *n_ps_* for population *p* of species *s* using a Binomial distribution

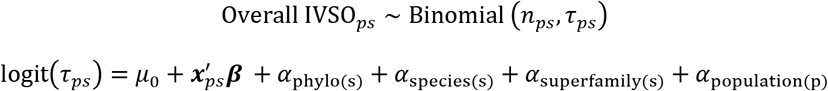

The probability *τ_ps_* predicts the proportion of social units expected to deviate from the primary social organization observed in the population or, equivalently, the probability of deviating for a randomly selected social unit.

Heterogeneous patterns of missing data were present for our ecological and life-history measures across species. As a consequence, a few populations lacked data for foraging style (1%), primary locomotion (2%), body size (2%), while many lacked data on the proportion of dietary reliance on fruits (18%), foliage (28%), seeds (60%), and animal protein (51%). Best practice for statistical estimation from a non-experimental dataset such as ours is to use some form of multiple imputation to account for non-random missingness across observations (38). Therefore, when seeking to assess the aggregate average effects of ecological predictors across species (Fig. 2A), we used the mice R package (42) to impute missing ecological and life-history values across predictors and obtain more reliable population statistics in our full model. However, for biological inferences about the effects of specific predictors (Fig 2B) and the overall ancestral state (Fig 3), we relied only on observed values taken from primary literature, excluding any rows containing missing data. Table S3 provides sample sizes and posterior estimates for all predictors in the full model with and without imputation, as well as in univariate models capturing the total effect of each predictor.

Ancestral state reconstructions are commonly carried out with intercept-only random effects models, in which the global model intercept is interpreted as the expected ancestral state after marginalizing over any species-level phylogenetic or stochastic effects; multiple regression models are then used separately to identify relevant selection pressures across the sample (36, 43). However, as the size and depth of the sampled phylogenetic tree grows, so too does the potential bias introduced into a reconstruction by unmeasured temporal trends and processes of non-random convergent evolution. These concerns are particularly acute for our dataset, which contains unbalanced samples across all major clades within the primate order and covers a span of approximately 51 million years. Therefore, we conducted our reconstruction of primate social organization in the context of a broader multiple regression model accounting for the effects of key social and ecological factors thought to be relevant to understanding the adaptive niche of ancestral primates, and which may also be associated with directional change in social organization and IVSO across extant primates. In particular, we assumed that ancestral primates were of relatively small body size (~30 g or - 2 SD z-score log mean body size), largely arboreal in their locomotion, and nocturnal in their activity pattern. We also adjusted our reconstruction for any biased sampling caused by differential research effort within primate superfamilies.

Given that our analyses were conducted in a fully Bayesian framework, we avoided the limitations of null-hypothesis testing (44), and instead relied on multiple sources of information provided by posterior distributions of model parameters and predictions. Median posterior estimates and median absolute deviations (MADs) were used to characterize the central tendency and relative dispersion of estimated effects, while 90% Bayesian credible intervals (CIs) and posterior probabilities of positive or negative effects (i.e. *p*+ or *p*−) were used to gauge uncertainty in the magnitude and direction of these effects (Table S4). Note that a 90% Bayesian CI excluding zero indicates greater than 0.95 posterior probability in support of a directional effect. These posterior probabilities *p*+ or *p*− directly quantify support for substantive rather than null hypotheses, so that values closer to 1 indicate greater support for the directional effect (+,−) and values closer to 0 indicate greater support for the opposite directional effect (−,+). All models were estimated in the Stan statistical programming language (45) using R (R Core Team 2020) and the brms package(46).

## Acknowledgments

We thank the CNRS, the University of Strasbourg and the University of Zurich. Comments by S.F. Dobson improved the significance statement.

## Supplementary Text

### Robustness checks

#### Phylogenetic process

A model assuming multivariate Brownian Motion for the phylogenetic effects generated nearly identical predictions as compared to a Gaussian Process phylogenetic model, which used a more flexible, nonparametric model of the phylogenetic effects analogous to an Orstein-Uhlenbeck process. In particular, differences in median predictions for ancestral social organizations and IVSOs were all < 0.05 between these models, suggesting that findings were robust to this analytic choice. We therefore opted for the simpler and more computationally efficient BM model.

#### Accounting for phylogenetic uncertainty

We needed to account for uncertainty in the phylogenetic relationships among taxa while making inferences from our model. For Bayesian inference, a straightforward solution is to random sample multiple phylogenetic covariance matrices during model estimation, capturing the uncertainty in the phylogenetic tree, and to combine across their posterior distributions for statistical inference. However, due to the computational costs of running a complex Bayesian model multiple times, it is desirable to select a sufficiently small number of random phylogenetic trees for which results are relatively robust to the addition of further samples. We tested this for our model by comparing naïve ancestral state inferences between models using a single or multiple random trees (N = 1, 5, 10). At N = 5 samples, we found that further addition of random trees (N = 10) did not meaningfully change inferences (posterior median differences < 0.01). We therefore pooled uncertainty across 5 random trees for all reported analyses.

---

**Table S1.** The different forms of social organization recorded in our study.

| **Social organization** | **Definition** |
| --- | --- |
| Solitary | Both resident adult solitary males and solitary females occur in the population.<br>All identified dispersers were excluded. Cases where only one sex occurred alone were considered as potential dispersal or alternative reproductive tactics and were not accounted as a solitary social organization. |
| Pair-living MF | One adult female and one adult male with or without dependent offspring. |
| Groups of one female and multiple males FMM | One adult female and multiple adult males. |
| Groups of one male and multiple females MFF | One adult male and multiple adult females. |
| Groups of multiple females and multiple males FFMM | Multiple adult females and multiple adult males. |
| Sex-specific group | Both multiple adult female groups and multiple adult male groups occur in the population.<br>Cases where only one sex-group occurred alone were considered as potential dispersal or alternative reproductive tactics and were not considered as a sex-specific social organization. |

**Table S2.** Predictors and from which database the information was used.

| Predictors | Definition | Database |
| --- | --- | --- |
| Habitat heterogeneity | Total number of habitats per species. | Based on our own primary literature search. |
| Habitat type | The type of habitat depending on whether it is open, closed or both.<br>Open: grassland, shrubland, rocky areas, savanna, desert.<br>Closed: forest, wetland, cave, woodland.<br>Both: artificial. | Based on our own primary literature search. |
| Body mass | Mean from the body mass of male and female; the standard deviation was used to account for measurement error. | (30, 31) |
| Diet | Percentage of the diet consisting of fruits and foliage throughout the entire year. | (30, 31) |
| Activity Pattern | Diurnal, nocturnal or cathemeral. | (30, 31) |
| Locomotion | Terrestrial, arboreal or both. | (30, 31) |
| Foraging strategy | Species that forage solitarily, in groups or both (solitarily and in groups). | Based on our own primary literature search. |

**Table S3.** Effect sizes for ecological predictors across models. Median posterior estimates are reported with posterior median absolute deviations (robust SDs) in parentheses. Estimates are taken from the full model including all ecological and life-history predictors with (N = 499 populations) or without imputation (N = 102 populations). Sample sizes are also reported in-line for total effect models with missing data, which only included fixed effects for research effort and the predictor of interest. Note that in a multinomial model, ecological effects on the total probability of each organization (e.g. as shown in Figure 2) can be estimated using a predictor’s effect on the difference in probability between solitary and the remaining social organizations (reported below).

| MF - Solitary |  |  |  |
| --- | --- | --- | --- |
| Effect size ($\beta$) and robust standard deviation (MAD) | | | |
| Predictor | Full model (imputation, N = 499 populations) | Full model (raw data, N = 102) | Total effect models (univariate) |
| Habitat heterogeneity | 0.12 (0.41) | 0.34 (0.56) | 0.16 (0.39) |
| Habitat category |  |  | N = 494 |
| -Closed v both | 0.75 (0.69) | 1.20 (0.88) | 0.75 (0.70) |
| -Open v both | -0.37 (0.91) | -0.05 (0.96) | -0.35 (0.88) |
| Foraging style |  |  | N = 496 |
| -Groups v solitary | 1.65 (0.76) | -0.12 (0.92) | 1.50 (0.72) |
| -Both v solitary | -0.38 (0.70) | 0.09 (0.91) | -0.30 (0.68) |
| Locomotion |  |  | N = 485 |
| -Both v Arb. | -0.18 (0.67) | -0.03 (0.82) | -0.36 (0.70) |
| -Terrest. v Arb. | 0.08 (0.90) | -0.43 (0.89) | -0.15 (0.80) |
| Activity pattern |  |  |  |
| -Cathem. v Noct. | -0.42 (0.87) | 0.16 (0.89) | 0.03 (0.84) |
| -Diurn. v Noct. | -0.67 (0.77) | -0.15 (0.93) | -0.80 (0.76) |
| Diet |  |  |  |
| -Fruit % | 0.47 (0.80) | 0.10 (0.93) | 0.43 (0.79); N = 409 |
| -Foliage % | 0.01 (0.81) | 0.31 (0.94) | 1.20 (0.79); N = 361 |
| -Seed % | -0.15 (0.89) | -0.12 (0.91) | -0.24 (0.92); N = 198 |
| -Animal % | 0.3 (0.87) | -0.38 (0.95) | 0.10 (0.86); N = 244 |
| Body size | -0.74 (0.35) | -1.70 (0.57) | -0.97 (0.36); N = 491 |
| MFF - Solitary |  |  |  |
| Effect size ($\beta$) and robust standard deviation (MAD) | | | |
| Predictor | Full model (imputation, N = 499 populations) | Full model (raw data, N = 102) | Total effect models (univariate) |
| Habitat heterogeneity | -0.06 (0.32) | 0.57 (0.48) | -0.07 (0.33) |
| Habitat category |  |  | N = 494 |
| -Closed v both | -0.42 (0.56) | -0.51 (0.74) | -0.45 (0.58) |
| -Open v both | 0.04 (0.71) | -0.59 (0.89) | 0.08 (0.74) |
| Foraging style |  |  | N = 496 |
| -Groups v solitary | 1.99 (0.74) | 0.19 (0.90) | 2.11 (0.74) |
| -Both v solitary | -1.41 (0.77) | -0.19 (0.91) | -1.38 (0.77) |
| Locomotion |  |  | N = 485 |
| -Both v Arb. | 0.36 (0.57) | -1.28 (0.81) | 0.41 (0.58) |
| -Terrest. v Arb. | -0.20 (0.70) | -0.56 (0.79) | -0.05 (0.71) |
| Activity pattern |  |  |  |
| -Cathem. v Noct. | 0.15 (0.86) | -0.20 (1.00) | 0.10 (0.86) |
| -Diurn. v Noct. | 1.22 (0.81) | 0.21 (0.98) | 1.52 (0.84) |
| Diet |  |  |  |
| -Fruit % | 0.03 (0.76) | 0.04 (0.90) | -0.33 (0.75); N = 409 |
| -Foliage % | 0.02 (0.72) | 0.20 (0.96) | 0.22 (0.76); N = 361 |
| -Seed % | 0.82 (0.86) | 0.00 (0.94) | -0.13 (0.88); N = 198 |
| -Animal % | 1.28 (0.82) | 0.60 (0.95) | 0.69 (0.86); N = 244 |
| Body size | 0.01 (0.36) | -0.23 (0.58) | 0.12 (0.36); N = 491 |
| FMM - solitary |  |  |  |
| Effect size ($\beta$) and robust standard deviation (MAD) | | | |
| Predictor | Full model (imputation, N = 499 populations) | Full model (raw data, N = 102) | Total effect models (univariate) |
| Habitat heterogeneity | -0.34 (0.46) | 0.11 (0.60) | -0.34 (0.47) |
| Habitat category |  |  | N = 494 |
| -Closed v both | 0.16 (0.65) | 0.56 (0.86) | 0.10 (0.66) |
| -Open v both | -0.34 (0.92) | -0.03 (1.00) | -0.29 (0.92) |
| Foraging style |  |  | N = 496 |
| -Groups v solitary | 2.62 (0.72) | 0.33 (0.93) | 2.81 (0.73) |
| -Both v solitary | -0.78 (0.70) | -0.33 (0.94) | -0.89 (0.71) |
| Locomotion |  |  | N = 485 |
| -Both v Arb. | 0.29 (0.62) | -0.08 (0.86) | 0.27 (0.66) |
| -Terrest. v Arb. | 0.08 (0.79) | -0.23 (0.97) | -0.02 (0.80) |
| Activity pattern |  |  |  |
| -Cathem. v Noct. | 0.89 (0.80) | -0.28 (0.95) | 1.03 (0.83) |
| -Diurn. v Noct. | 0.85 (0.77) | 0.29 (0.92) | 0.99 (0.76) |
| Diet |  |  |  |
| -Fruit % | -0.05 (0.73) | -0.51 (0.90) | -0.28 (0.75); N = 409 |
| -Foliage % | -0.12 (0.75) | 0.08 (0.96) | 0.25 (0.76); N = 361 |
| -Seed % | 0.91 (0.86) | 0.63 (0.96) | 0.58 (0.86); N = 198 |
| -Animal % | 0.06 (0.83) | -0.12 (1.00) | 0.17 (0.83); N = 244 |
| Body size | -0.33 (0.29) | -1.21 (0.59) | -0.23 (0.33); N = 491 |
| FFMM - Solitary |  |  |  |
| Effect size ($\beta$) and robust standard deviation (MAD) | | | |
| Predictor | Full model (imputation, N = 499 populations) | Full model (raw data, N = 102) | Total effect models (univariate) |
| Habitat heterogeneity | 0.17 (0.36) | 0.22 (0.59) | 0.23 (0.36) |
| Habitat category |  |  | N = 494 |
| -Closed v both | -0.29 (0.56) | -0.66 (0.73) | -0.36 (0.54) |
| -Open v both | 0.65 (0.70) | 0.60 (0.91) | 0.61 (0.73) |
| Foraging style |  |  | N = 496 |
| -Groups v solitary | 2.19 (0.77) | 1.10 (0.86) | 2.23 (0.80) |
| -Both v solitary | -1.37 (0.76) | -1.11 (0.91) | -1.37 (0.79) |
| Locomotion |  |  | N = 485 |
| -Both v Arb. | 0.73 (0.59) | 0.18 (0.78) | 0.77 (0.59) |
| -Terrest. v Arb. | 0.17 (0.71) | 0.99 (0.79) | 0.47 (0.71) |
| Activity pattern |  |  |  |
| -Cathem. v Noct. | 0.43 (0.85) | 0.35 (0.96) | 0.47 (0.88) |
| -Diurn. v Noct. | 0.94 (0.80) | -0.40 (0.95) | 1.23 (0.82) |
| Diet |  |  |  |
| -Fruit % | -0.61 (0.74) | -0.15 (0.94) | -0.58 (0.75); N = 409 |
| -Foliage % | -0.99 (0.71) | -0.30 (1.01) | -0.57 (0.79); N = 361 |
| -Seed % | 0.25 (0.78) | 0.06 (0.94) | 0.14 (0.91); N = 198 |
| -Animal % | -0.34 (0.84) | -0.24 (0.96) | 0.17 (0.87); N = 244 |
| Body size | 0.33 (0.38) | 0.38 (0.59) | 0.36 (0.40); N = 491 |
| Total IVSO |  |  |  |
| Effect size ($\beta$) and robust standard deviation (MAD) | | | |
| Predictor |  |  |  |

|  | Full model (imputation, N<br>= 499 populations) | Full model (raw data,<br>N = 102) | Total effect models<br>(univariate) |
| --- | --- | --- | --- |
| Habitat heterogeneity | -0.14 (0.18) | -0.10 (0.29) | -0.08 (0.18) |
| Habitat category |  |  | N = 494 |
| -Closed v both | 0.17 (0.35) | 0.87 (0.57) | 0.09 (0.33) |
| -Open v both | -0.20 (0.50) | -0.72 (0.86) | -0.01 (0.51) |
| Foraging style |  |  | N = 496 |
| -Groups v solitary | -0.39 (0.56) | -0.73 (0.83) | -0.17 (0.50) |
| -Both v solitary | 0.59 (0.55) | 0.71 (0.85) | 0.58 (0.57) |
| Locomotion |  |  | N = 485 |
| -Both v Arb. | -0.37 (0.30) | -0.56 (0.59) | -0.26 (0.30) |
| -Terrest. v Arb. | 0.44 (0.42) | 0.37 (0.64) | 0.37 (0.38) |
| Activity pattern |  |  |  |
| -Cathem. v Noct. | 0.81 (0.68) | -0.47 (0.94) | 0.41 (0.65) |
| -Diurn. v Noct. | 0.93 (0.55) | 0.48 (0.94) | 0.74 (0.52) |
| Diet |  |  |  |
| -Fruit % | -0.83 (0.57) | -0.49 (0.84) | -1.16 (0.48); N = 409 |
| -Foliage % | 0.11 (0.55) | -0.22 (0.84) | 0.70 (0.51); N = 361 |
| -Seed % | 0.39 (0.79) | -0.15 (0.83) | 0.11 (0.73); N = 198 |
| -Animal % | 1.04 (0.70) | 0.36 (0.94) | 0.55 (0.67); N = 244 |
| Body size | 0.11 (0.16) | 0.20 (0.35) | 0.11 (0.16); N = 491 |

**Table S4.** Correlation of social organization and IVSO across species Species-level correlations. Median posterior correlations are shown for the phylogenetic and species residual effects on the relative probability of pair and group living in comparison to solitary social organization. Posterior probabilities in support of a positive correlation *p*+ shown in parentheses. Posterior probabilities lower than 0.95 and greater than 0.05 provide weak evidence of a positive or negative effect, respectively. Overall, little evidence was found for species-level correlations between specific forms of social organization and overall IVSO, with most posterior correlations centered near zero with relatively high uncertainty. No clear associations were observed for residual species differences in social organization or IVSO. Therefore, to enhance computational efficiency, we simplified our statistical models by assuming independent evolution of social organization and IVSO across the phylogenetic tree. However, specific forms of social organization did tend to coevolve within lineages, as indicated by moderate positive correlations between the relative probability of MF and MFF (median *r* = 0.52, *p*_+_ = 0.99) and FMM (median *r* = 0.67, *p*_+_ = 1.00) in comparison to solitary living. The relative probability of MFF was also positively correlated with FMM (median *r* = 0.67, *p*_+_ = 1.00) and FFMM (median *r* = 0.63, *p*_+_ = 1.00), as well as the relative probability of FMM and FFMM (median *r* = 0.58, *p*_+_ = 0.99). See Table S4 for all results.

**Phylogenetic correlation**
|  | Solitary/MF | Solitary/MFF | Solitary/FMM | Solitary/FFMM | Overall IVSO |
| --- | --- | --- | --- | --- | --- |
| Solitary/MF | 1 |  |  |  |  |
| Solitary/MFF | 0.52 (0.99) | 1 |  |  |  |
| Solitary/FMM | 0.67 (1.00) | 0.67 (1.00) | 1 |  |  |
| Solitary/FFMM | -0.00 (0.49) | 0.63 (1.00) | 0.58 (0.99) | 1 |  |
| Overall IVSO | 0.13 (0.73) | 0.16 (0.80) | 0.32 (0.94) | -0.17 (0.33) | 1 |

**Species residual correlation**
|  | Solitary/MF | Solitary/MFF | Solitary/FMM | Solitary/FFMM | Overall IVSO |
| --- | --- | --- | --- | --- | --- |
| Solitary/MF | 1 |  |  |  |  |
| Solitary/MFF | 0.06 (0.56) | 1 |  |  |  |
| Solitary/FMM | 0.05 (0.55) | -0.00 (0.50) | 1 |  |  |
| Solitary/FFMM | -0.03 (0.47) | 0.01 (0.51) | 0.07 (0.53) | 1 |  |
| Overall IVSO | 0.03 (0.53) | 0.11 (0.53) | 0.09 (0.59) | -0.17 (0.24) | 1 |

## Reference

1. P. M. Kappeler, L. Pozzi, Evolutionary transitions toward pair living in nonhuman primates as stepping stones toward more complex societies. Science Advances 5, eaay1276 (2019).

2. D. Lukas, T. Clutton-Brock, Evolution of social monogamy in primates is not consistently associated with male infanticide. Proceedings of the National Academy of Sciences 111, E1674 (2014).

3. S. Shultz, C. Opie, Q. D. Atkinson, Stepwise evolution of stable sociality in primates. Nature 479, 219–222 (2011).

4. C. Opie, Q. D. Atkinson, R. I. M. Dunbar, S. Shultz, Male infanticide leads to social monogamy in primates. Proceedings of the National Academy of Sciences 110, 13328–13332 (2013).

5. D. Lukas, T. H. Clutton-Brock, The evolution of social monogamy in mammals. Science 341, 526–530 (2013).

6. P. Agnani, C. Kauffmann, L. D. Hayes, C. Schradin, Intra-specific variation in social organization of Strepsirrhines. American Journal of Primatology 80, e22758 (2018).

7. P. M. Kappeler, Lemur behaviour informs the evolution of social monogamy. Trends in Ecology & Evolution 29, 591–593 (2014).

8. P. M. Kappeler, C. P. v. Schaik, Evolution of primate social systems. Int J Primatol 23, 707–740 (2002).

9. P. M. Kappeler, A framework for studying social complexity. Behavioral Ecology and Sociobiology 73, 13 (2019).

10. M. Huck, A. Di Fiore, E. Fernandez-Duque, Of apples and oranges? The evolution of “monogamy” in non-human primates. Frontiers in Ecology and Evolution 7 (2020).

11. E. Fernandez-Duque, M. Huck, S. Van Belle, A. Di Fiore, The evolution of pair-living, sexual monogamy, and cooperative infant care: Insights from research on wild owl monkeys, titis, sakis, and tamarins. American Journal of Physical Anthropology 171, 118–173 (2020).

12. P. A. Garber, L. M. Porter, J. Spross, A. Di Fiore, Tamarins: Insights into monogamous and non-monogamous single female social and breeding systems. American Journal of Primatology 78, 298–314 (2016).

13. C. Kvarnemo, Why do some animals mate with one partner rather than many? A review of causes and consequences of monogamy. Biological Reviews 93, 1795–1812 (2018).

14. S. R. Tecot, B. Singletary, E. Eadie, Why “monogamy” isn’t good enough. American Journal of Primatology 78, 340–354 (2016).

15. A. Cohas, D. Allainé, Social structure influences extra-pair paternity in socially monogamous mammals. Biology letters 5, 313–316 (2009).

16. L. Brouwer, S. C. Griffith, Extra-pair paternity in birds. Mol. Ecol. 28, 4864–4882 (2019).

17. C. A. Chapman, A. Corriveau, V. A. M. Schoof, D. Twinomugisha, K. Valenta, Long-term simian research sites: significance for theory and conservation. Journal of Mammalogy 98, 652–660 (2017).

18. J. H. Crook, J. S. Gartlan, Evolution of primate societies. Nature 210, 1200–1203 (1966).

19. T. Clutton-Brock, C. Janson, Primate socioecology at the crossroads: Past, present, and future. Evolutionary Anthropology: Issues, News, and Reviews 21, 136–150 (2012).

20. R. I. M. Dunbar, “Socio-ecological systems” in Primate Social Systems, R. I. M. Dunbar, Ed. (Cornell University Press, New York, 1988), pp. 262–291.

21. C. Schradin, L. D. Hayes, N. Pillay, C. Bertelsmeier, The evolution of intraspecific variation in social organization. Ethology 124, 527–536 (2018).

22. X. Ni et al., The oldest known primate skeleton and early haplorhine evolution. Nature 498, 60–64 (2013).

23. M. A. O’Leary et al., The placental mammal ancestor and the post–K-Pg radiation of placentals. Science 339, 662–667 (2013).

24. L. V. Valen, R. E. Sloan, The earliest primates. Science 150, 743–745 (1965).

25. P. M. Kappeler et al., Long-term field studies of lemurs, lorises, and tarsiers. Journal of Mammalogy 98, 661–669 (2017).

26. A. V. Jaeggi, M. I. Miles, M. Festa-Bianchet, C. Schradin, L. D. Hayes, Variable social organization is ubiquitous in Artiodactyla and probably evolved from pair-living ancestors. Proceedings of the Royal Society B: Biological Sciences 287, 20200035 (2020).

27. M. Valomy, L. D. Hayes, C. Schradin, Social organization in Eulipotyphla: evidence for a social shrew. Biology Letters 11, doi:10.1098/rsbl.2015.0825 (2015).

28. C.-A. Olivier, A. V. Jaeggi, L. D. Hayes, C. Schradin, Revisiting the components of Macroscelidea social systems: Evidence for variable social organization, including pair-living, but not for a monogamous mating system. Ethology 128, 383–394 (2022).

29. D. R. Rubenstein, P. Abbot, “Opportunities for comparative social evolution” in Comparative Social Evolution, D. R. Rubenstein, P. Abbot, Eds. (Cambridge University Press, Cambridge, 2017), DOI: 10.1017/9781107338319.002, pp. 427–452.

30. L. Makuya, C.-A. Olivier, C. Schradin, Field studies need to report essential information on social organisation – independent of the study focus. Ethology 128, 268–274 (2022).

31. C. C. Grueter et al., Multilevel Organisation of Animal Sociality. Trends in Ecology & Evolution 35, 834–847 (2020).

32. H. Kummer, Social organization of hamadryas baboons (Karger, Basel, 1968).

33. F. Aureli et al., Fission-Fusion Dynamics New Research Frameworks. Current Anthropology 49 (2008).

34. C. Galán-Acedo, V. Arroyo-Rodríguez, E. Andresen, R. Arasa-Gisbert, Ecological traits of the world’s primates. Scientific Data 6, 55 (2019).

35. N. Rowe, M. Myers, “All The World’s Primates”. (2017), 10.1002/9781119179313.wbprim0086.

36. J. D. Hadfield, S. Nakagawa, General quantitative genetic methods for comparative biology: phylogenies, taxonomies and multi-trait models for continuous and categorical characters. Journal of Evolutionary Biology 23, 494–508 (2010).

37. T. E. Currie, A. Meade, “Keeping Yourself Updated: Bayesian Approaches in Phylogenetic Comparative Methods with a Focus on Markov Chain Models of Discrete Character Evolution” in Modern phylogenetic comparative methods and their application in evolutionary biology, L. Z. Garamszeg, Ed. (Springer, 2014), pp. 263–286.

38. R. McElreath, Statistical rethinking: A Bayesian course with examples in R and Stan (2nd editio) (CRC Press, 2020).

39. N. S. Upham, J. A. Esselstyn, W. Jetz, Inferring the mammal tree: Species-level sets of phylogenies for questions in ecology, evolution, and conservation. PLOS Biology 17, e3000494 (2019).

40. J. Koster, R. McElreath, Multinomial analysis of behavior: statistical methods. Behavioral Ecology and Sociobiology 71, 138 (2017).

41. N. P. Lemoine, Moving beyond noninformative priors: why and how to choose weakly informative priors in Bayesian analyses. Oikos 128, 912–928 (2019).

42. S. van Buuren, K. Groothuis-Oudshoorn, mice: Multivariate Imputation by Chained Equations in R. Journal of Statistical Software 45, 1 – 67 (2011).

43. T. Garland, Jr., A. R. Ives, Using the Past to Predict the Present: Confidence Intervals for Regression Equations in Phylogenetic Comparative Methods. Am Nat 155, 346–364 (2000).

44. B. B. McShane, D. Gal, A. Gelman, C. Robert, J. L. Tackett, Abandon Statistical Significance. The American Statistician 73, 235–245 (2019).

45. B. Carpenter et al., Stan: A Probabilistic Programming Language. Journal of Statistical Software 76, 1 – 32 (2017).

46. P.-C. Bürkner, brms: An R Package for Bayesian Multilevel Models Using Stan. 2017 80, 28 (2017).

